# Predicting Spiking Activity from Scalp EEG

**DOI:** 10.1101/2025.08.06.668998

**Authors:** Dixit Sharma, Bart Krekelberg

**Affiliations:** Center for Molecular and Behavioral Neuroscience, Rutgers University, Newark, NJ, 07102

## Abstract

Despite decades of electroencephalography (EEG) research, the relationship between EEG and underlying spiking dynamics remains unclear. This limits our ability to infer local neural dynamics reflected in intracranial signals from EEG, a critical step to bridge electrophysiological findings across species and to develop non-invasive brain-machine interfaces (BMIs). We recorded spiking activity from a 32-channel floating microarray permanently implanted in parafoveal V1 and scalp-EEG in a male macaque monkey. While the animal fixated, the screen flickered at different temporal frequencies to induce steady-state visual evoked potentials (SSVEP). We analyzed the relationship between the V1 multi-unit spiking activity envelope (MUAe) and EEG frequency bands to predict MUAe at each time point from EEG. We extracted instantaneous spectrotemporal features of the EEG signal, including phase, amplitude, and phase-amplitude coupling of its frequency bands. Although the relationship between these features and the V1 MUAe was complex and frequency-dependent, it was reliably predictive of the MUAe. Specifically, in a multivariate linear regression predicting MUAe from EEG, each EEG feature (phase, amplitude, coupling) contributed to model predictions. In addition, we found that MUAe predictions were better in shallow than deep cortical layers, and that the phase of stimulus frequency further improved MUAe predictions. Our study shows that a comprehensive account of spectrotemporal features of non-invasive EEG can inform about underlying spiking activity, which is beyond what is available when the amplitude or phase of the EEG signal is considered separately. This demonstrates the richness of the EEG signal and its complex relationship with neural spiking activity and suggests that using more comprehensive spectrotemporal signatures could improve BMI applications.

## INTRODUCTION

Electroencephalography (EEG) is a widely used non-invasive method for measuring neuronal activity in humans, valued for its cost-effectiveness, portability, and high temporal resolution. Despite its limited spatial resolution compared to imaging techniques like fMRI, EEG’s ability to capture neural dynamics at high spectrotemporal resolution has made it an essential tool for diagnosing neurological disorders (Fisher et al., 2005; Jiao et al., 2023), investigating the neural correlates of cognitive processes (Biasiucci et al., 2019; Friston, 2009), and controlling non-invasive brain-computer interfaces (Birbaumer et al., 2008).

There is a broad consensus that EEG signals primarily originate from the synchronized electrical activity of cortical neurons, specifically the (excitatory or inhibitory) postsynaptic potentials of pyramidal neurons oriented perpendicular to the cortical surface (Nunez & Srinivasan, 2006). Their parallel (or open-field) arrangement allows for spatial summation of postsynaptic potentials, resulting in strong extracellular dipole fields measured as EEG signals. However, in some cases, neurons in close-field arrangements, such as stellate cells, can also contribute to EEG activity (Tenke et al., 1993; see Nunez & Srinivasan, 2006 for a detailed review). Although the physics underlying EEG signals is well understood, research exploring how the spectrotemporal features of EEG relate to underlying cortical signals, such as spiking activity and local field potentials (LFP), remains limited.

A long-term objective of linking EEG with underlying cortical signals measured invasively is to develop tools that can estimate time-precise local cortical dynamics, reflected in neuronal spikes and LFPs, using non-invasive EEG signals. However, due to signal averaging and various noise factors that contribute to EEG, the low signal-to-noise ratio (SNR) makes this a challenging goal. Although complete realization may not be feasible in the short term, incremental advancements can help bridge the gap between invasive and non-invasive signals, and thus between animal and human neurophysiology research. Moreover, better understanding of EEG’s spectrotemporal features could improve clinical biomarkers and support the development of non-invasive brain–machine interfaces (BMIs) and closed-loop neuromodulation systems (Lu et al., 2021).

One promising approach to narrowing the gap between invasive and non-invasive recordings is to record both signals simultaneously in animals engaged in cognitive tasks. A few studies have used this approach to better understand the relationship between cortical signals and EEG. For example, Whittingstall & Logothetis (2009) found that multi-unit spiking activity in macaque V1 was weakly correlated with bone-surface EEG amplitude (r = -0.12), but showed a stronger relationship with EEG delta phase (2-4 Hz) and gamma power (30-100 Hz).

Interestingly, this frequency-dependent relationship can even be exploited to estimate V1 spiking activity (Whittingstall & Logothetis, 2009). Synchronization among cortical signals also impacts EEG: Snyder et al. (2015) reported a non-linear relationship between spike synchronization in V4 and EEG alpha-band power, with higher alpha power observed during both low and high spike synchronization compared to moderate levels. The relationship between EEG and cortical signals also seems to depend upon the provided visual stimulus (Snyder & Smith, 2015).

With a high-level goal of estimating cortical dynamics non-invasively, we build on previous studies to develop a comprehensive framework that permits estimations of time-precise spike dynamics in macaque V1 using non-invasive EEG. Our approach incorporates two major advancements over prior studies. First, to overcome the challenge of low SNR of EEG signals, we used a periodic flickering visual stimulus to elicit high SNR responses, known as steady state visual evoked potentials (SSVEPs). SSVEP responses are less susceptible to artifacts and exhibit high SNR (Norcia et al., 2015), and hence, are widely used in human EEG, including applications in investigating higher cognitive functions through frequency tagging (Vialatte et al., 2010; Zhu et al., 2010). It is important to note that SSVEP responses depend on the stimulus frequency (Alonso-Prieto et al., 2013), with frequencies within alpha and beta range (10–25 Hz) eliciting the strongest responses (Ladouce et al., 2022).

Second, leveraging the high temporal resolution of EEG, we developed a comprehensive predictive model to estimate V1 spiking activity using non-invasive EEG. Specifically, we used amplitude, phase, and phase-amplitude coupling features of EEG bands to linearly model V1 spiking activity at each time point. Given that neural activity patterns may vary across cortical layers (Mendoza-Halliday et al., 2024; Xing et al., 2012), we also evaluated the model’s performance according to different cortical layers.

Our prediction model significantly exceeded chance performance, with markedly higher accuracy during SSVEP stimulus trials. Specifically, the model demonstrated better prediction accuracy for trials involving SSVEP stimuli compared to non-SSVEP trials, and superficial cortical layers were predicted more accurately than deeper layers. Given the non-invasive nature of EEG in our study, this approach has the potential for direct translation to human EEG applications.

## METHODS

### Subject, Recording Procedures, and Apparatus

The subject was an adult male rhesus macaque (Macaca mulatta) weighing approximately 18 kg at the time of neural and behavioral data recordings. All experimental procedures were performed following the NIH Guide for the Care and Use of Laboratory Animals and the ARVO Statement for the Use of Animals in Ophthalmic and Vision Research, with approval from the Rutgers University Animal Care and Use Committee. Surgical implantation was conducted under fully aseptic conditions while the animal was under ketamine-induced isoflurane anesthesia. Ibuprofen and morphine analgesia were given as required post-surgery.

Neurophysiological signals were recorded while the animal engaged in a fixation task. Intracranial spiking activity was recorded using a 32-channel floating microarray, permanently implanted in the parafoveal region of the left hemisphere V1. The individual electrodes on the microarray (1.2 x 3.4 mm²) were spaced at 400 μm both horizontally and vertically. Electrodes were distributed across four depths, with eight electrodes at each depth; the electrode length ranged from 0.6 to 1.5 mm relative to the dura surface, and the array was inserted perpendicular to the dura surface. EEG signals were concurrently recorded using three non-invasive scalp Ag/AgCl electrodes placed above the occipital lobe (see Fig. 1b), with online reference and ground EEG electrodes placed above the left and right frontal lobe, respectively.

**Fig. 1.**
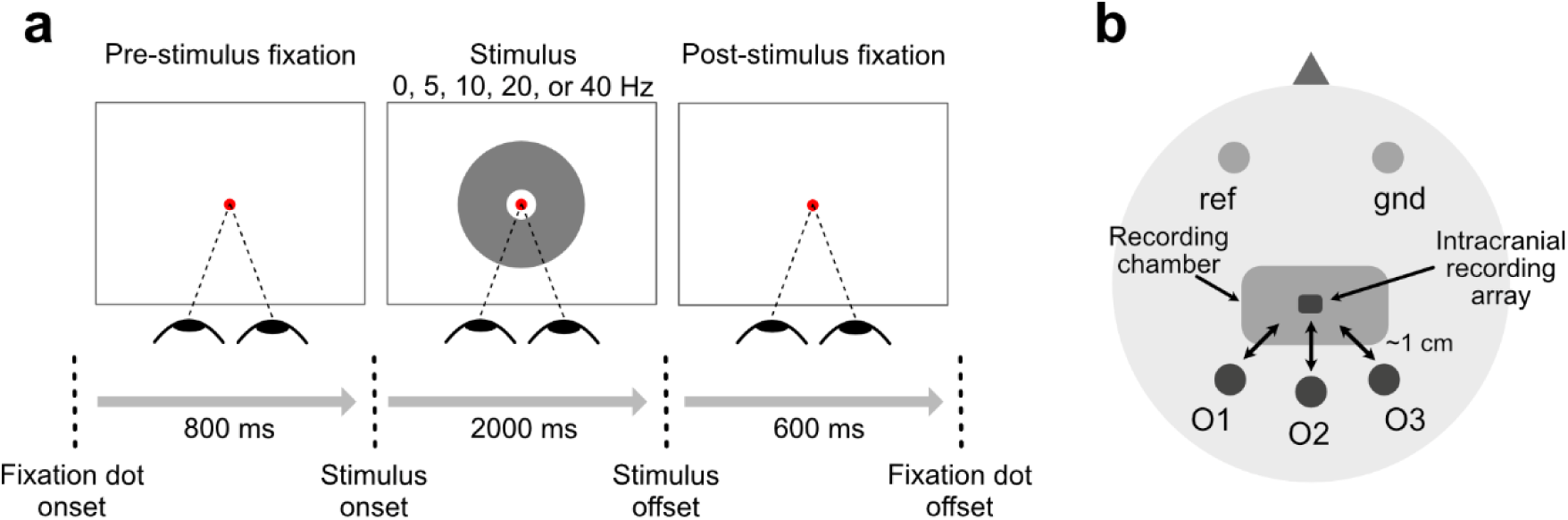
Behavioral task and EEG montage. (a) Task sequence (not shown to scale), in which monkeys fixated on a central dot continuously in the pre-stimulus (800 ms), stimulus (2000 ms), and post-stimulus (600 ms) phase of the task to receive a juice reward. During the stimulus phase, the screen flickers at one of the five stimulus frequencies (0, 5, 10, 20, or 40 Hz). (b) Schematic of EEG electrode montage. Each dark circle indicates the position of EEG electrodes on the occipital cortex and their associated labels (O1, O2, O3). The reference (ref) and ground (gnd) electrodes were placed over the frontal cortex and are marked in light gray circles. The round-edged rectangle indicates the position of the surgically implanted chamber used to protect the connectors of the intracranial recording array. The bidirectional arrows indicate the approximate distance (1 cm) between intracranial and EEG electrodes.

During the recording sessions, the animal was seated comfortably in a standard primate chair without head restraint, approximately 57 cm away from a 120 Hz cathode-ray tube monitor. Eye movements were continuously monitored using an infrared video eye tracking system (EyeLink 1000; SR Research, Ottawa, Ontario, Canada) at a sampling rate of 1000 Hz. Stimulus presentation and eye tracking were controlled via our custom software Neurostim (Krekelberg et al., 2022) and the Psychophysics Toolbox (Brainard, 1997; Pelli, 1997) in MATLAB (MathWorks Inc., Natick, MA). Neurophysiological, eye movement, and behavioral data were synchronously acquired with the Ripple system. Data analysis was performed using custom MATLAB scripts alongside standard analysis tools.

### Behavioral Task

The animal performed a dot-fixation task (Fig. 1a), in which it was required to maintain its gaze within 1.5° of a central fixation dot for the entire trial duration (3400 ms). A trial was considered successful when the animal fixated throughout three consecutive phases of the trial: pre-stimulus (800 ms), stimulus (2000 ms), and post-stimulus (600 ms).

During the pre-stimulus phase, the monkey initiated a trial by fixating on the central dot against a gray background for 800 ms, triggering the onset of the stimulus phase. In the stimulus phase, a donut-shaped stimulus (radius = 8.9 cm) centered around the fixation dot was displayed over the background. The donut flickered at one of five pseudorandomly selected temporal frequencies: 0, 5, 10, 20, or 40 Hz, referred to as trial conditions. In the 0 Hz condition, the donut appeared and remained stationary on the screen throughout the stimulus period, serving as an open-eye spontaneous condition without flicker. The flickering stimuli (5, 10, 20, and 40 Hz) were used to evoke steady-state visual evoked potentials (SSVEP), a response known to produce high signal-to-noise ratios (SNR) in both EEG and intracranial signals (Norcia et al., 2015).

During the post-stimulus phase, the background reverted to gray. A liquid reward was delivered contingent upon the animal maintaining fixation throughout all three phases. A 250 ms intertrial interval, during which no visual stimuli were present, followed each trial. Trials in which fixation was broken were aborted, and data from these trials were excluded from subsequent analyses.

### Neurophysiological recordings and signals

Neural signals from the primary visual cortex (V1) and non-invasive EEG were simultaneously recorded in the monkey during task performance. The primary goal of our acquisition setup was to record both intracranial (from V1) and extracranial (EEG) signals concurrently, while minimizing skull alteration to facilitate human-like, non-invasive EEG recordings. Therefore, we trained the animal to perform the task without a head-restraining device and kept the skull intact except at the site of chronic intracranial microarray implantation.

The intracranial electrode array implanted in V1 recorded the unfiltered wideband neural signals at a sampling rate of 30 kHz. We extracted multi-unit activity envelope (MUAe) from wideband signals offline using a procedure developed by (Supèr & Roelfsema, 2005). This procedure involves the following steps sequentially: 1) band-pass filtering the wideband signal (500-5000 Hz), 2) rectifying the filtered signal, 3) applying the Hilbert transform to the rectified signal, and 4) downsampling to 1 kHz.

EEG was recorded at a sampling rate of 1 kHz with an online low-pass IIR filter of 500 Hz. Since the animal was not head-restrained, conductive adhesive gel (Ten20; Weaver and Company, Aurora, Colorado, USA) was applied to ensure stable electrode contact throughout the recording sessions. Before each session, the animal’s scalp was shaved and cleaned with saline and alcohol to achieve low impedance. Electrodes with impedance below 20 kOhm at the beginning of each recording session were included in subsequent analysis.

We applied bidirectional FIR digital filters of order 1000 on the recorded EEG signal to extract full-band and band-limited signals. We first removed power line noise using bidirectional bandpass FIR filters (60, 120, and 180 Hz). For full-band EEG, we then applied high-pass (1 Hz) and low-pass (250 Hz) filters. For band-limited signals we applied bandpass filters according to frequency band of interest: delta (2-4 Hz), theta (4-8 Hz), alpha (8-15), beta (15-30 Hz), low-gamma (30-60 Hz), gamma (60-120 Hz), and high-gamma (120-200 Hz). The full-band and band-limited signals were segmented into trials and aligned to the stimulus onset.

### Electrode and trial selection

To identify functional MUAe electrodes from our chronic implant, we visually inspected the power spectrum of each electrode across multiple example sessions, focusing on spectral peaks at stimulus frequencies. Power spectra were computed over the 500-1500 ms window after stimulus onset using the Fast Fourier Transform (FFT). All cortical electrodes displayed prominent peaks at the stimulus frequencies. Hence, we included all 32 cortical electrodes in our analysis.

Next, for each session, we assessed whether the signal measured on EEG electrode remained responsive to task stimuli. This step was important because our animal was not head-restricted, which sometimes led to excessive head movement. Such movement could cause EEG electrodes to shift or loosen, resulting in poor contact and compromised signal quality within the session. To quantify the signal quality of each EEG electrode in each session, we calculated the signal-to-noise ratio (SNR), defined as the ratio of evoked power at the stimulus frequency to the average power at the two neighboring frequencies. Only EEG electrodes with an SNR > 1.5 for at least one stimulus condition in a session were retained for further analysis. Applying these criteria resulted in 31 EEG electrodes across 14 sessions, yielding a total of 992 EEG-MUAe electrode pairs used in our analysis.

Artifact trials were identified separately using EEG and LFP signals. For LFP, trials were marked as artifacts if they met any of the following criteria: (1) high synchronization relative to other trials (z > 4) across more than 25% of electrodes; (2) outlier signals relative to other trials (z > 4) in at least two electrodes; or (3) LFP magnitude exceeding 500 μV in more than two electrodes. For EEG, artifact trials were identified using either of the following criteria: (1) outlier signal relative to other trials (z > 4) at any EEG electrode; or (2) EEG magnitude exceeding 100 μV for over 10 ms in any electrode. Trials identified by either method were excluded from further analysis. This resulted in 4318 trials used for further analysis, averaging 308 trials per session (SD = 105).

### Data Analysis

We focused our analysis on finding relationships between MUAe and EEG with the goal of developing a comprehensive linear model to predict V1 spike dynamics using EEG. For all our analyses below we used a 400-1800 ms timeseries signal following stimulus onset, either using the single-trial timeseries (“single-trial”) or timeseries signals averaged across trials (“trial-averaged”). All analyses were performed in MATLAB (R2019b).

#### Correlation across time

To quantify the relationship between EEG and MUAe signals, we calculated Pearson’s correlation coefficients between their timeseries, separately for each stimulus condition. These correlations were computed for both full-band and band-limited EEG across all EEG-MUAe electrode pairs.

#### Phase-amplitude coupling

We used the Modulation Index (MI, Tort et al., 2008) to quantify the coupling between the phase of EEG signals and the amplitude of both EEG and MUAe. To calculate MI, phase values were binned into 18 intervals of 20° from -180° to 180°. The corresponding EEG amplitude or MUAe was averaged for each phase bin to construct the phase-amplitude distribution. Following the methodology of Hülsemann et al. (2019), we then computed the MI as the Kullback-Leibler divergence between the observed phase-amplitude distribution and a uniform distribution. We chose MI over alternatives such as phase-locking value or mean vector length because MI can detect biphasic coupling and is less susceptible to confounding factors like signal-to-noise ratio and data length (Hülsemann et al., 2019).

#### Multivariate regression analysis

We applied L1-regularized multivariate linear regression (LASSO) to predict the instantaneous MUAe at each time point from EEG spectrotemporal features. These features included the analytic amplitude and the instantaneous phase of seven frequency bands, as well as their pairwise couplings.. The regression model is expressed in Wilkinson’s notation as:

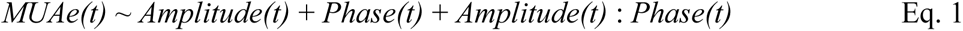

where *Amplitude* and *Phase* denote the time series of EEG amplitudes and phases across the seven frequency bands, and *t* indicates time sample. Due to circular nature of phase variable, we transformed phases via sine and cosine functions to enable linear modeling of phase effects. The phase-related beta coefficients were derived by calculating the Euclidean norm of the sine and cosine beta estimates. All predictors were z-scored across samples prior to model fitting.

Based on our phase-amplitude coupling analysis (Fig. 5), we included interaction terms (Amplitude : Phase) between the amplitudes of beta, low-gamma, gamma, and high-gamma bands and the phases of the delta, theta, and alpha bands. Separate models were constructed for single-trial and trial-averaged signals.

Model performance was evaluated using a 5-fold cross-validation procedure. In each iteration, time samples from 4 folds were used to train the model with LASSO regression, and the trained coefficients were applied to predict MUAe in the held-out test samples. The average model performance across all five folds was reported.

To determine the optimal regularization parameter (λ) for the LASSO regression, we performed a nested 5-fold cross-validation on the training data, following L1 norm criterion (Hastie et al., 2015). During this process, a set of λ values was evaluated, and the λ that minimized the average mean squared error across the inner folds was selected as the optimal regularization parameter. The coefficients corresponding to this λ were then estimated using the entire training set and used to predict MUAe in the test data during each fold. This approach ensured an unbiased and robust estimate of model accuracy on unseen data.

#### Statistical analysis

Since we recorded from multiple electrodes simultaneously – 32 MUAe electrodes and 3 EEG electrodes – the measurements are likely not fully independent. Measurements from electrodes within the same session may be partially correlated, which means the usual assumptions of independence in many statistical tests do not strictly hold in this dataset. To address this, we used linear mixed-effect models, with random intercept effects for each EEG- MUAe electrode pair to account for potential dependencies between electrode pairs and a separate random intercept effect for each session to account for repeated measurements within a session.

Specifically, we used MATLAB’s *fitlme.m* function to implement mixed-effect models, which allowed us to evaluate the statistical measures of EEG–MUAe correlation coefficients and model performance metrics. To account for the effect of repeated measurements on the degrees of freedom and corresponding p-values, we calculated the degrees of freedom using the Satterthwaite approximation method, ensuring more accurate statistical values.

## RESULTS

The primary objective of this study was to predict V1 spike dynamics from non-invasive EEG recordings at millisecond resolution. We simultaneously recorded multi-unit activity envelopes (MUAe) from V1 and scalp EEG over the occipital cortex in a macaque monkey (Fig. 1b). Neural data were collected during a fixation task in which the screen flickered at one of five temporal frequencies (0, 5, 10, 20, and 40 Hz) per trial (Fig. 1a), with the frequency selected pseudorandomly on each trial. These stimuli evoked high signal-to-noise ratio steady-state visual potentials (SSVEPs; Norcia et al., 2015). For all analyses, we used signals from 400 to 1800 ms post-stimulus onset to minimize transient effects related to stimulus onset and offset.

### Correlations between MUAe and full-band EEG are weak and stimulus-dependent

To evaluate a direct relationship between EEG and MUAe we first calculated Pearson’s correlation between full-band EEG and MUAe timeseries signals averaged across trials (i.e., trial-averaged timeseries). Fig. 2 shows example EEG and MUAe timeseries signals and their corresponding spectrum. In this example, correlation coefficient for spontaneous (0 Hz) and flickering (10 Hz) condition is -0.004 and 0.16, respectively, indicating that SSVEP responses evoked by flickering stimuli are better correlated between EEG and MUAe. Fig. 3 shows distribution of correlation coefficients across all EEG-MUAe pairs for each stimulus condition. Similar to the pattern seen in example signals, correlations for 0 Hz condition was non-significant (r = -0.008 ± 0.010, p = 0.37) and 10 Hz condition show positive correlations (r = 0.06 ± 0.01, p = 3.0e-6).

**Fig. 2.**
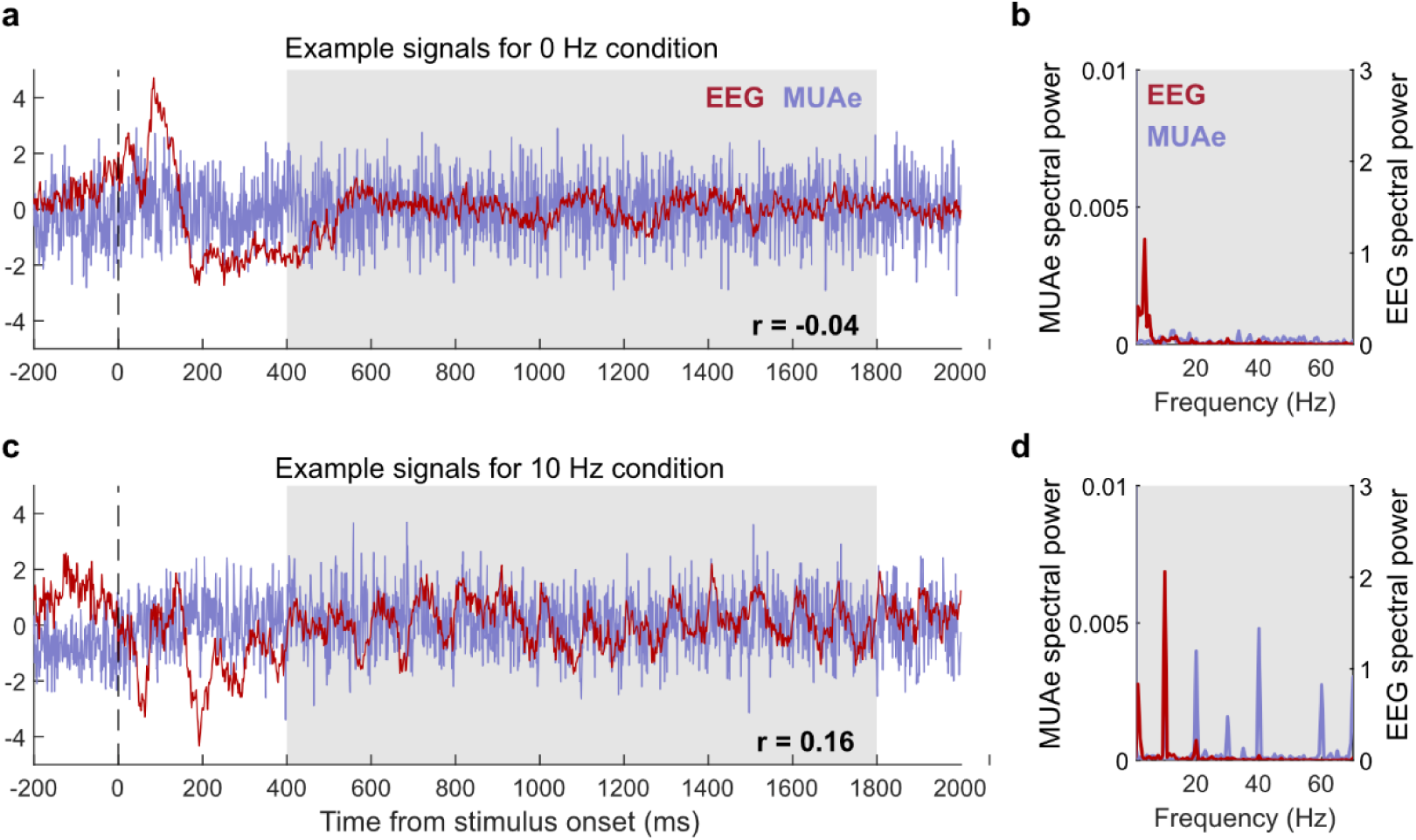
Neural signals and corresponding spectral representations from representative electrodes. (a) Neural signals from representative electrodes for spontaneous (0 Hz) condition, showing time series MUAe (blue) and EEG (red) signal averaged across trials; a clear onset response can be seen in EEG signal. (b) Power spectra of the signals from panel (a) in the 400-1800 ms time window (indicate by the shaded gray region). (c) Same as (a), but for trials with a flickering (10 Hz) stimulus condition. (d) Same as (b), but for signals shown in shaded region of panel (c).

**Fig. 3.**
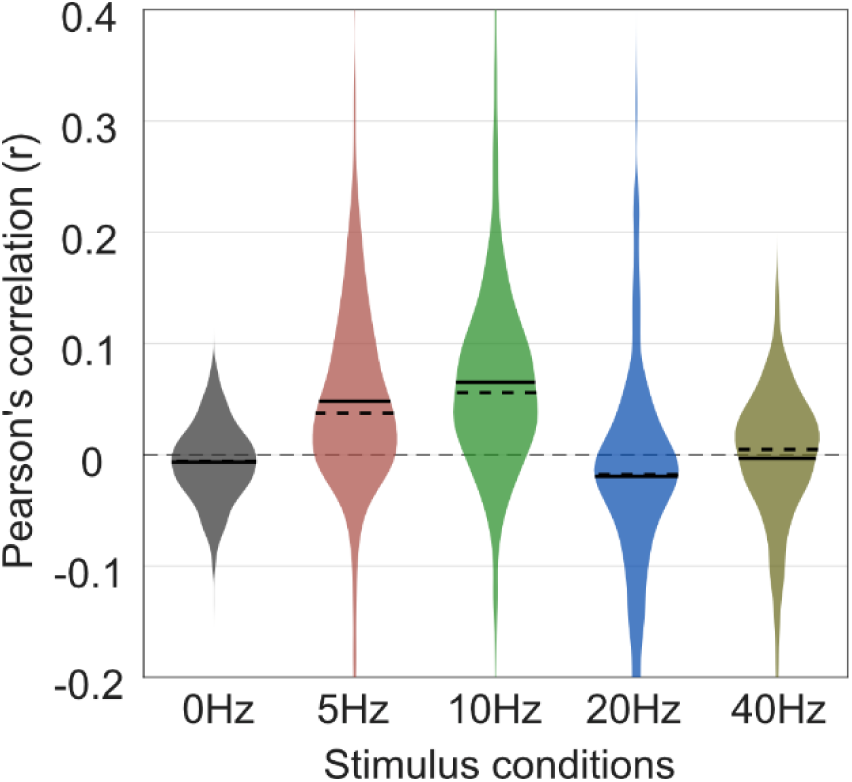
Pearson’s correlations across time between full-band EEG and MUAe for each stimulus condition (0, 5, 10, 20, and 40 Hz). Each violin indicates correlation distribution across EEG-MUAe electrode pairs recorded over sessions.

EEG-MUAe relationship further varied depending on frequency of stimulus conditions. Whereas 5 and 10 Hz conditions showed positive correlations (5 Hz, r = 0.05 ± 0.01, p = 1.0e-4; 10 Hz, r = 0.06 ± 0.01, p = 3.0e-6), correlation for 20Hz was slightly negative (r = -0.02 ± 0.01, p = 0.03) and non-significant for 40 Hz (r = -0.005 ± 0.010, p = 0.57). These results indicate that the strength of the EEG–MUAe relationship depends strongly upon the stimulus frequency, corroborating earlier findings (Snyder & Smith, 2015).

### Relationship between EEG and V1 spiking depends on frequency bands in EEG

We then investigated how the phase and amplitude of EEG bands – delta (2–4 Hz), theta (4–8 Hz), alpha (8–15 Hz), beta (15–30 Hz), low-gamma (30–60 Hz), gamma (60–120 Hz), and high-gamma (120–200 Hz) – relate to V1 MUAe. The correlation analysis revealed that the relationships between band-limited EEG timeseries and MUAe were not uniform across frequency bands nor limited to a single band, but partially distributed across bands and depended on the stimulus frequency (Fig. 4a).

**Fig. 4.**
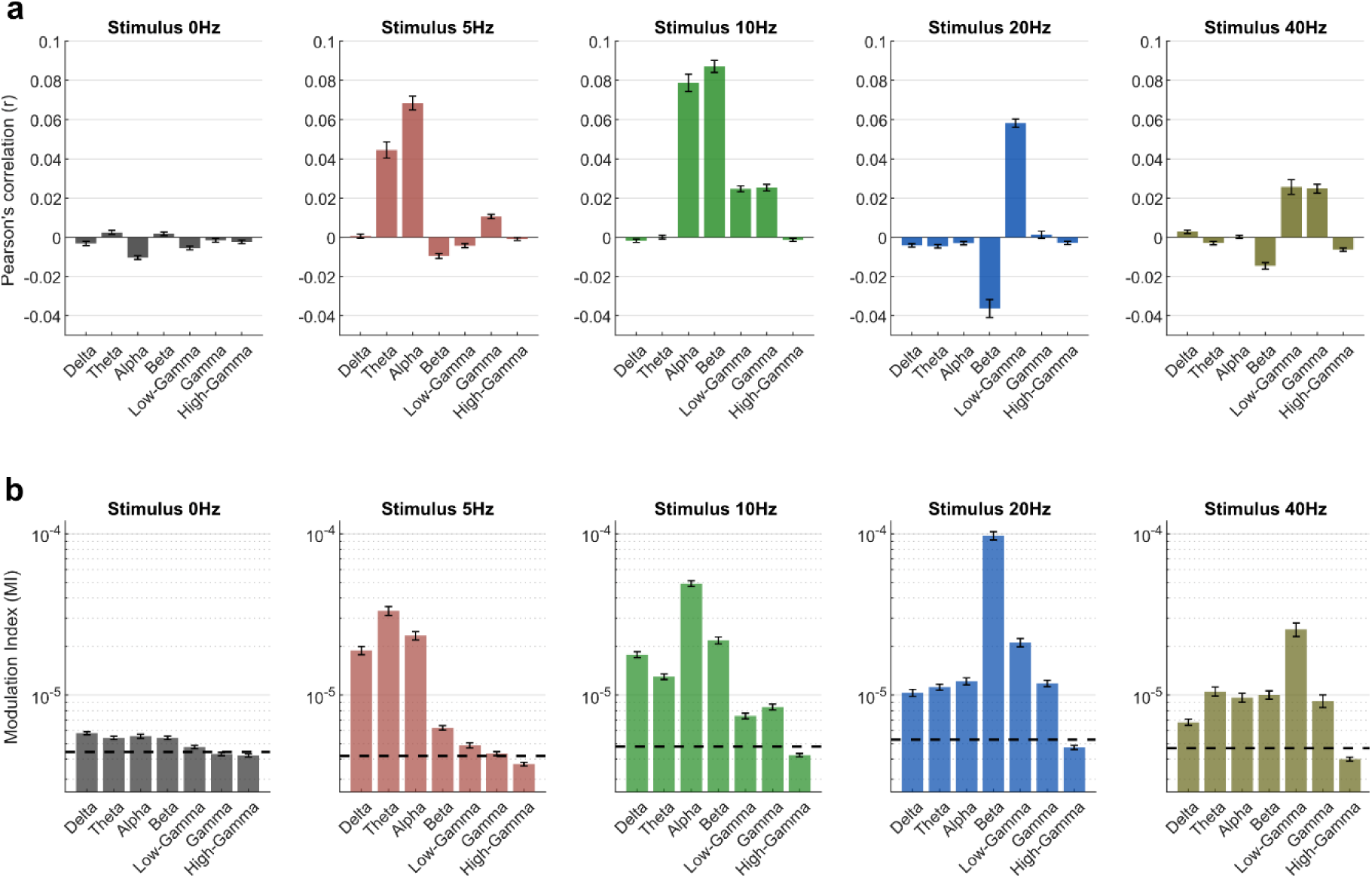
Relationship between MUAe and EEG-bands. (a) Pearson’s correlations across time between MUAe and EEG-bands (delta, theta, alpha, beta, low-gamma, gamma, and high-gamma). Each bar shows the mean correlation for EEG-MUAe electrode pairs recorded over sessions, shown separately for each stimulus condition (0, 5, 10, 20, and 40 Hz). (b) Modulation index (MI) quantifying phase-amplitude coupling between MUAe and phase of EEG-bands. Each bar shows the mean MI for EEG-MUAe electrode pairs, shown separately for each stimulus condition (0, 5, 10, 20, and 40 Hz). Dashed lines indicate mean MI calculated for MUAe and shuffled EEG phase bins iterated 1000 times. Error bars indicate SEM.

In the non-flicker spontaneous (0 Hz) condition, correlations across all bands were weak and near zero, with slight negative trends observed in delta (r = –0.003 ± 0.002, p = 0.046), alpha (r = –0.01 ± 0.002, p = 7.1e-8), and low-gamma (r = –0.005 ± 0.002, p = 9.4e-4). During flickering stimuli at 5, 10, 20, and 40 Hz, the significant correlations tended to occur in bands resonant with the stimulus frequency or its harmonics.

For the 5 Hz stimulus, positive correlations were observed in the theta (r = 0.043 ± 0.005, p = 1.5e-7) and alpha (r = 0.067 ± 0.005, p = 8.8e-11) along with some negative trend in beta (r = –0.012 ± 0.005, p=0.037) bands. The 10 Hz condition showed higher positive correlations, particularly in the alpha (r = 0.083 ± 0.007, p = 3.3e-10), beta (r = 0.091 ± 0.007, p = 6.9e-11), low-gamma (r = 0.027 ± 0.007, p = 6.3e-4) and gamma (r = 0.028 ± 0.007, p = 5.0e-4) bands. Interestingly, in the 20 Hz condition, two neighboring bands showed significant correlations with opposite trends: whereas beta band showed negative (r = -0.039 ± 0.005, p = 3.3e-7), low-gamma showed positive correlation (r = 0.056 ± 0.005, p = 6.2e-10). Finally, the 40 Hz stimulus elicited positive correlations in the low-gamma (r = 0.021 ± 0.006, p = 0.005) and gamma (r = 0.020 ± 0.006, p = 0.008) bands, but negative in beta band (r = -0.020 ± 0.006, p = 0.008). Overall, these correlation results indicated that all the frequency bands can be informative about EEG-MUAe relationships, depending on the type of stimulus used in the task.

We next evaluated if phase of EEG bands are related to MUAe activity in any band-specific way. To do so, we calculated phase-amplitude coupling (PAC) using Modulation Index (MI; Tort et al., 2008) between the phase of EEG bands and MUAe. Similar to the correlation results above, PAC between EEG bands and MUAe depended upon the stimulus conditions. For spontaneous (0 Hz) condition, couplings are generally weak with only slightly above the shuffled value for delta to beta bands. In contrast, flickering stimuli (i.e., 5, 10, 20, & 40 Hz stimuli) showed higher MI for certain bands relative to shuffled data, according the stimulus condition (Fig. 4b). Specifically, the 5 Hz stimulus showed increased MI for delta to beta bands, while the 10, 20, and 40 Hz stimuli showed high MI for all bands except high-gamma.

These results highlight the frequency-specific nature of EEG–MUAe interactions, emphasizing the importance of examining both phase and amplitude of each EEG band when studying EEG-spike relationships, further extending previous findings (Whittingstall & Logothetis, 2009).

### Amplitude and phase of EEG bands predict V1 spike dynamics

Leveraging observations from the correlation and PAC analysis, we implemented a cross-validated, L1-regularized, multivariate regression model to estimate MUAe at each time point from EEG features, including amplitudes, phases, and interaction terms of EEG bands (Eq. 1).

These interaction terms accounted for PAC between EEG amplitude and phase, a phenomenon observed in several human studies (Canolty & Knight, 2010). We hypothesized that PAC between EEG bands might provide unique information beyond what is available from amplitude or phase alone, thereby potentially enhancing MUAe prediction accuracy (Canolty & Knight, 2010; Fell & Axmacher, 2011).

We first assessed which pairs showed strong PAC by calculating MI for each phase-amplitude pair. As shown in Fig. 5 (white box), the phase of low-frequency bands (delta, theta, and alpha) coupled with the amplitude of high-frequency bands (beta, low-gamma, gamma, and high-gamma). Based on this observation, we included 12 interaction terms in our model: pairing the phase of three bands (delta, theta, and alpha) with the amplitude of four bands (beta, low-gamma, gamma, and high-gamma) (Eq. 1).

The model was 5-fold cross-validated across time samples. Pearson’s correlations (r) between the estimated MUAe and original MUAe for the left-out samples measured model performance, and the average of correlation across folds is reported.

**Fig. 5.**
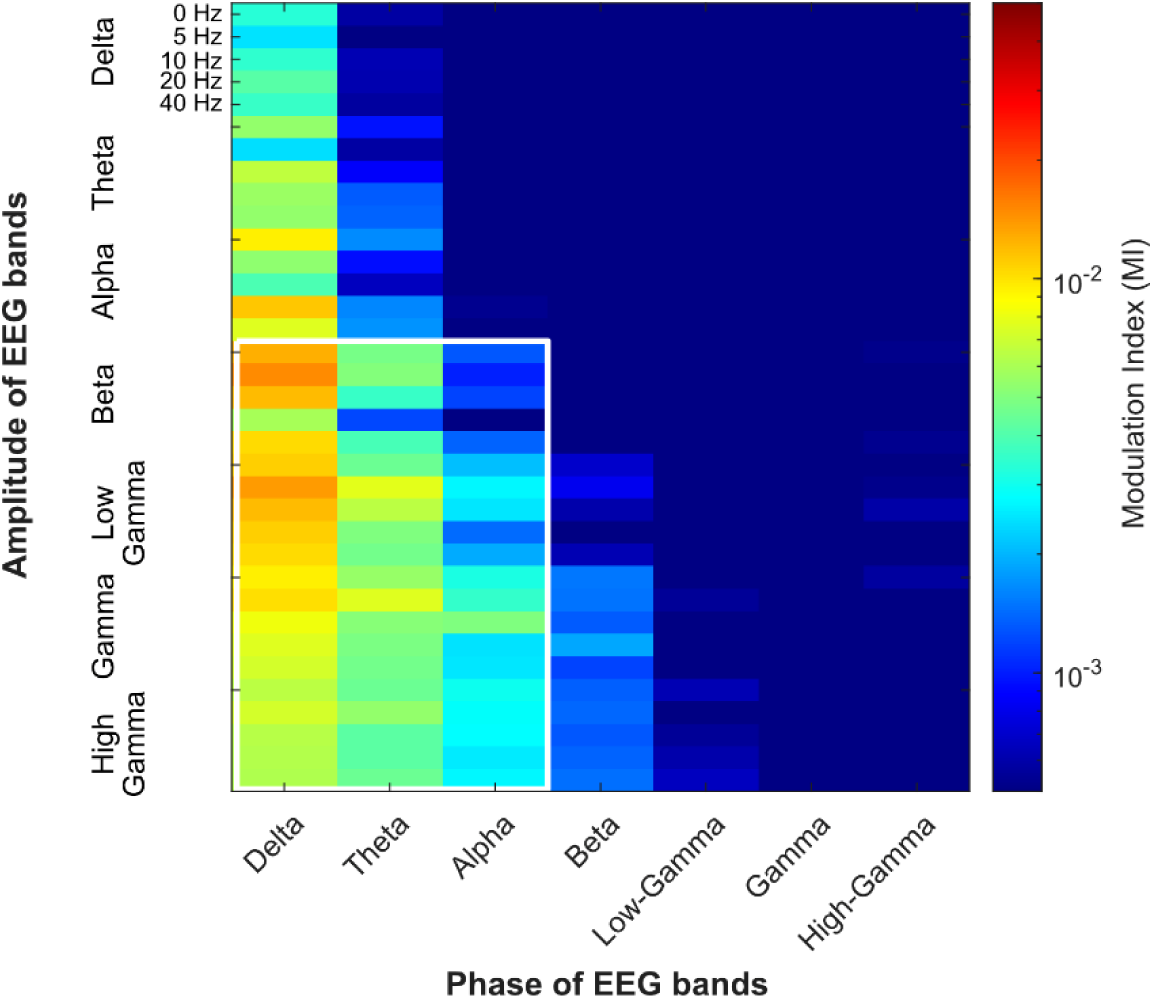
Coupling between phase and amplitude of EEG bands. The color axis indicates the coupling between phases (x-axis) and amplitudes of EEG bands (y-axis) for each stimulus condition (0, 5, 10, 20, and 40 Hz) quantified by the Modulation Index (MI). The white rectangle indicates phase-amplitude coupling pairs we used in the model predicting MUAe using spectrotemporal features of EEG signal (Eq. 1).

Fig. 6a shows the distribution of correlations across EEG-MUAe electrode pairs and sessions for trial-averaged signals. The model performed well for all flickering stimuli (5, 10, 20, and 40 Hz) but not for non-flickering 0 Hz stimuli (r = 0.02 ± 0.022, p = 0.35). The best performance was observed for the 10 Hz stimulus (r = 0.30 ± 0.022, p = 4.0e-11), followed by 20 Hz (r = 0.26 ± 0.022, p = 4.8e-10), 5 Hz (r = 0.26 ± 0.022, p = 5.1e-10), and 40 Hz (r = 0.17 ± 0.022, p = 5.1e-7).

**Fig. 6.**
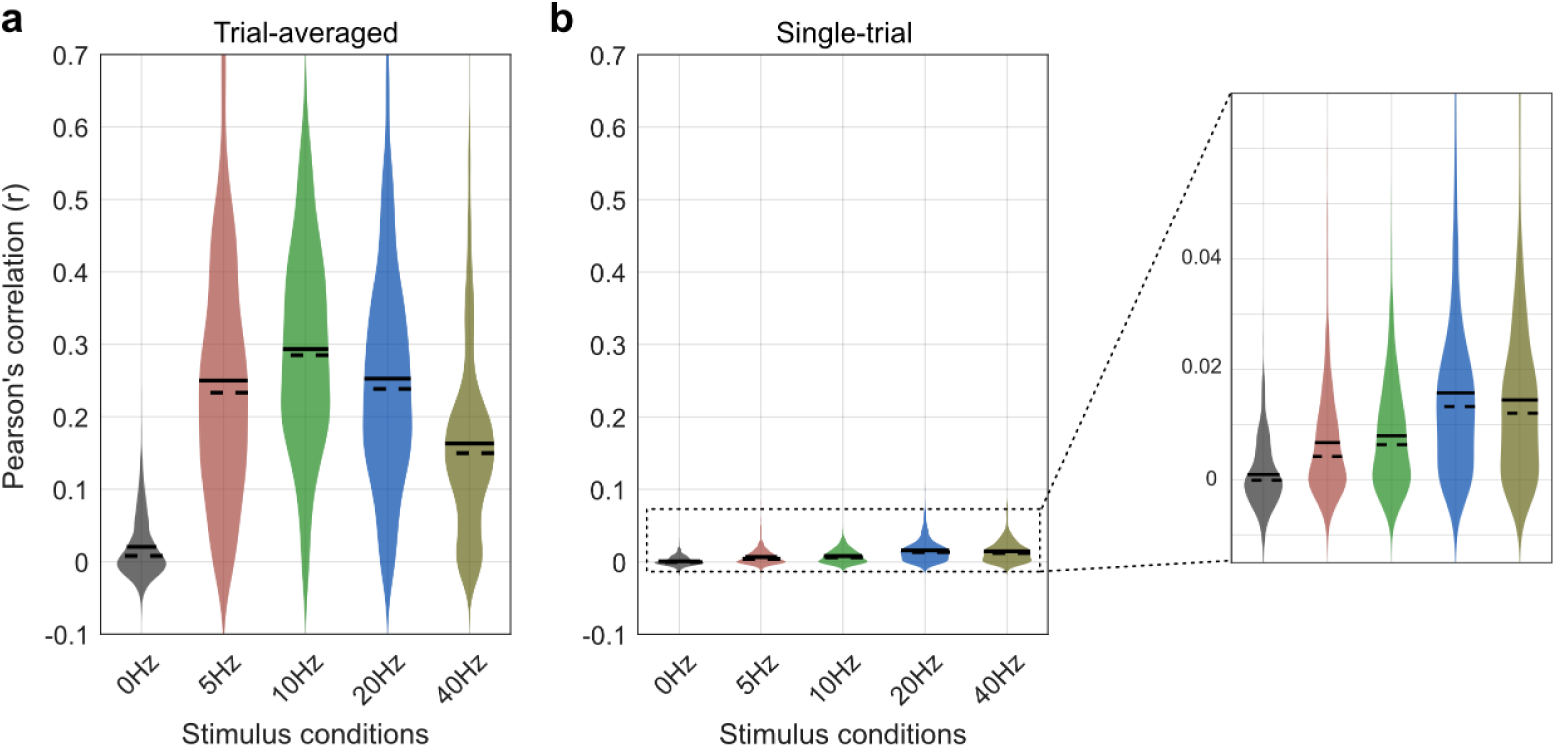
Model prediction accuracy quantified using Pearson’s correlations between timeseries of original and predicted MUAe. (a) Correlation between original and predicted trial-averaged MUAe signals. Each violin indicates the distribution of correlation between the original and predicted MUAe for a stimulus condition across all EEG-MUAe electrode pairs recorded over sessions. (b) Same as (a) but for correlations calculated using single-trial MUAe signals. The inset shows an enlarged y-axis.

Compared to trial-averaged signals, model performance was worse for trial-level signals (Fig. 6b). At the trial-level, the model performed worst for 0 Hz stimulus (r=0.001± 0.001, p = 0.55) but better for flickering stimuli (5 Hz, r = 0.006 ± 0.001, p = 2.0e-5; 10 Hz, r = 0.008 ± 0.001, p= 1.6e-6; 20 Hz, r = 0.016 ± 0.001, p = 4.9e-12; 40 Hz, r = 0.014 ± 0.001, p = 2.8e-11). A closer look at the correlations revealed that the 20 and 40 Hz stimuli elicited higher correlations than 5 and 10 Hz stimuli (see inset in Fig. 6b).

To compare the performance of our model relative to direct EEG–MUAe correlations (Fig. 3), we calculated the difference between the correlation coefficient of the EEG–MUAe pair and that of the estimated versus actual MUAe. Our model showed better correlations than direct EEG–MUAe in all stimulus conditions, for both trial-averaged (0 Hz, Δr = 0.027, p = 8.3e-6; 5 Hz, Δr = 0.213, p < 1.0e-30; 10 Hz, Δr = 0.239, p < 1.0e-30; 20 Hz, Δr = 0.278, p < 1.0e-30; 40 Hz, Δr = 0.170, p < 1.0e-30) and single-trial signals (0 Hz, Δr = 0.001, p = 0.065; 5 Hz, Δr = 0.005, p < 2.7e-14; 10 Hz, Δr = 0.005, p < 9.9e-16; 20 Hz, Δr = 0.016, p < 1.0e-30; 40 Hz, Δr = 0.012, p < 1.0e-30).

To better understand the relative contribution of amplitude, phase, and interaction terms used in the model, we compared the absolute values of the regression coefficients (betas) for these (z-scored) predictor variables. We first grouped absolute betas of all predictor variables in three categories: amplitude of seven bands, phase of seven bands, and coupling indicated by interaction terms of the model (Supp. Fig. 2), and compared them using linear-mixed effect model (see Methods).

Comparing amplitude and phase categories for trial-averaged signals revealed significant differences between them: whereas the spontaneous 0 Hz condition showed greater contribution of amplitude variables (difference in absolute beta coefficient, Δb = 0.001, p = 1.0e-12), flickering conditions showed greater contribution of phase variables (5 Hz, Δb = -0.002, p = 1.6e-4; 10 Hz, Δb = -0.005, p = 7.4e-9; 20 Hz, Δb = -0.009, p = 3.6e-20; 40 Hz, Δb = -0.003, p = 1.8e-10). Similarly, comparison of amplitude and coupling categories showed greater betas for amplitude for 0 Hz condition (Δb = 0.0004, p = 1.4e-7) and inverse for flickering conditions (5 Hz, Δb = -0.006, p = 1.5e-15; 10 Hz, Δb = -0.009, p = 4.8e-14; 20 Hz, Δb = -0.001, p = 7.8e-5; 40 Hz, Δb = -0.002, p = 2.5e-7). Overall, these results suggested that for trial-averaged EEG-to- MUAe estimations, amplitude variables contribute more for spontaneous conditions, but for flickering conditions, phase and coupling variables contribute more.

Interestingly, for single-trial estimations, phase variables always contributed more than amplitude variables for all stimulus conditions (0 Hz, Δb = -0.001, p = 1.2e-35; 5 Hz, Δb = - 0.003, p = 1.87e-27; 10 Hz, Δb = -0.003, p = 3.63e-30; 20 Hz, Δb = -0.005, p = 2.95e-28; 40 Hz, Δb = -0.004, p = 1.82e-23). Similar was the case for the amplitude and coupling comparison (0 Hz, Δb = -0.0007, p = 6.9e-31; 5 Hz, Δb = -0.0007, p = 6.7e-15; 10 Hz, Δb = -0.0009, p = 2.1e-17; 20 Hz, Δb = -0.0009, p = 8.0e-12; 40 Hz, Δb = -0.0009, p = 9.3e-11). These results suggested that phase and coupling are almost always more informative for EEG-to-MUAe predictions than amplitude variables.

### Phase of stimulus frequency improves EEG-to-spike predictions

As mentioned above, Whittingstall & Logothetis (2009) found that the phase of EEG delta band (2–4 Hz) can help predict spiking activity in V1 during naturalistic movie viewing tasks. Mazzoni et al. (2010) explained this by suggesting that V1 is adapted to naturalistic sensory stimuli, which tend to induce oscillations within the delta frequency range. They argued that this is why EEG delta phase provides valuable information about V1 spiking activity.

Building on this idea, Mazzoni et al. (2010) hypothesized that any stimulus or stimulation causing V1 activity to oscillate in a regular, periodic manner could improve the prediction of V1 spiking. Specifically, they proposed that the phase of stimulus frequencies could serve as a key predictor of V1 spiking activity.

To test this hypothesis, we used EEG and V1 MUAe data recorded during flickering visual stimuli and evaluated if phase of a flickering stimulus improved EEG-to-MUAe predictions. In particular, we updated our prediction model (Eq. 1) by adding stimulus phase as a predictor variable, and compared the explained variance (adjusted r^2^) of models with and without the stimulus phase. We found that using stimulus phase as an additional predictor in the model significantly improve the model performance, particularly at a single-trial level (Fig. 7). For trial-averaged estimations, only 40 Hz condition showed significant improvement (difference in adjusted r², Δr² = 0.087, p = 1.3e-10); the model performance was not different for any other stimulus condition (5 Hz, Δr² = 0.002, p = 0.77; 10 Hz, Δr² = 0.003, p = 0.70; 20 Hz, Δr² = - 0.007, p = 0.29). For trial-level estimations, model with stimulus phase performed better for all stimulus conditions (5 Hz, Δr² = 0.0012, p = 0.0018; 10 Hz, Δr² = 0.0015, p = 0.0003; 20 Hz, Δr² = 0.0011, p = 0.0044; 40 Hz, Δr² = 0.0026, p = 7.3e-8), supporting the hypothesis proposed by Mazzoni et al. (2010).

**Fig. 7.**
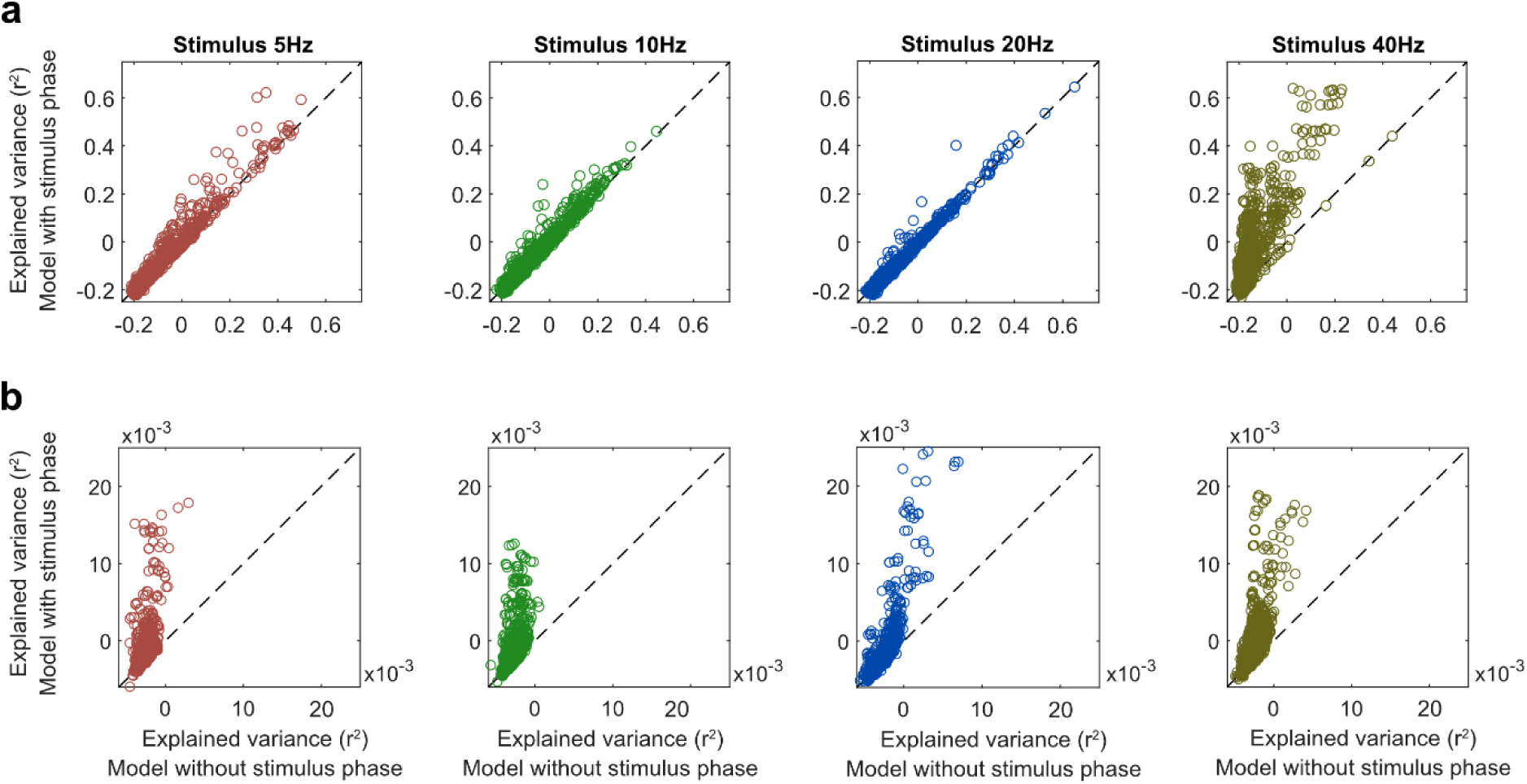
Scatter plots comparing prediction models with and without stimulus phase as a predictor. The x-axis shows the explained variance of the prediction model excluding the stimulus phase as a predictor, and y-axis shows the prediction for the model that includes stimulus phase as a predictor (Eq. 1). (a) Predictions using trial-averaged MUAe and EEG signals. (b) Predictions using single-trial MUAe and EEG signals. Each sub-panel indicates a condition with a flickering, sinusoidal stimulus (5, 10, 20, 40 Hz).

### Spiking activity predictions are better at superficial than deep layers

Multiple studies have shown that spiking and LFP activity show distinct patterns across cortical layers (Bastos et al., 2018; Mendoza-Halliday et al., 2024; Mitzdorf, 1985). This suggests that our EEG-to-MUAe model prediction performance may also vary across cortical layers in V1. To investigate this, we compared model prediction accuracy between MUAe recorded at different cortical depths.

We recorded MUAe from 32 electrodes placed at four equidistant depths, putatively spanning superficial (2/3) to deep cortical layers (4, 5, 6) in V1. To directly compare the model performance at superficial versus deep layers, we grouped the electrodes into two categories: shallow (covering layers 2/3) and deep (covering layers 4-6). The laminar placement of these electrodes was confirmed using the methodology developed by Mendoza-Halliday et al. (2024). We then evaluated the model’s prediction accuracy separately for these groups.

As shown in Fig. 8, MUAe predictions at superficial layers are generally higher than deep layers, particularly for flickering stimuli. For trial-averaged signals, all flickering conditions showed significant higher model performance at shallow than deep layers (5 Hz, Δr = 0.063, p = 0.0003; 10 Hz, Δr = 0.049, p = 0.0026; 20 Hz, Δr = 0.036, p = 0.017; 40 Hz, Δr = 0.027, p = 0.052), which was not seen for spontaneous 0 Hz condition (Δr = -0.002, p = 0.55). Similar trend was observed for single-trial signals for both flickering and spontaneous conditions (0 Hz, Δr = 0.000, p = 0.49; 5 Hz, Δr = 0.002, p = 0.079; 10 Hz, Δr = 0.003, p = 0.0051; 20 Hz, Δr = 0.006, p = 3.2e-6; 40 Hz, Δr = 0.004, p = 0.0019). These results suggest that EEG predicts spiking activity more accurately in the superficial layers, supporting the idea that superficial layers contribute more to the EEG signal than deeper layers.

**Fig. 8.**
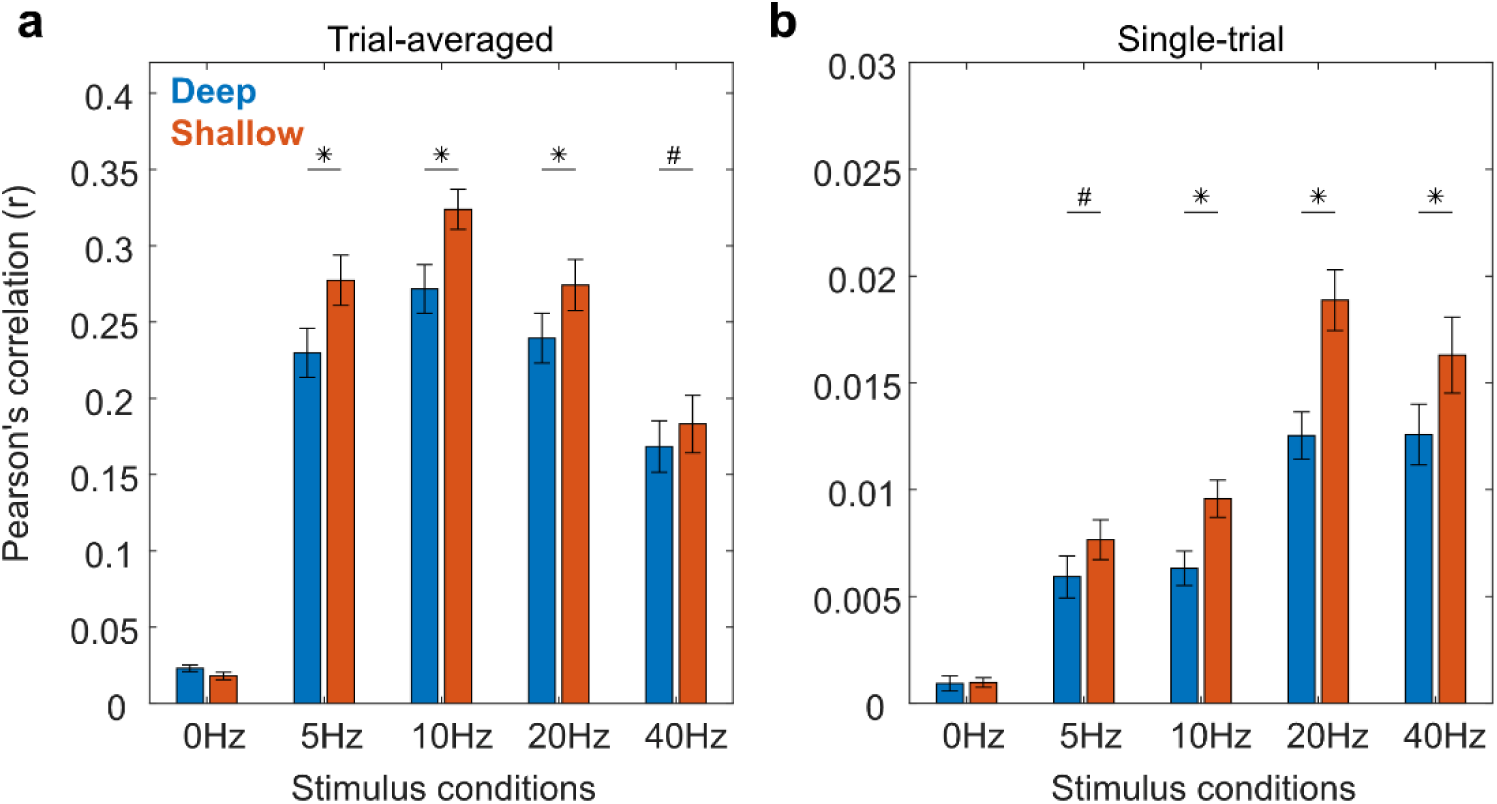
Bar plots comparing EEG-to-MUAe model predictions at deep (blue) and shallow (orange) cortical depths of V1 recording sites. Significant differences in model predictions between deep and shallow MUAe electrode sites are indicate by * (p < 0.05) and # (p < 0.10). (a) Prediction using trial-averaged EEG and MUAe signals. (b) Predictions using single-trial EEG and MUAe signals.

## DISCUSSION

### Summary

Neuronal spiking activity is a local signal reflecting both stimulus and cognitive information dynamically at millisecond resolutions, making it a primary focus of investigation in neuroscience. However, due to its invasive recording nature, this signal is typically studied in animals. In contrast, EEG, which is non-invasive and typically recorded in humans, offers sub-second temporal resolution but is constrained by poor SNR and spatial resolution. The relationship between these two signals is not well established, limiting the potential to estimate neural dynamics non-invasively with high temporal precision.

With the high-level goal of estimating temporally precise neural activity non-invasively, this study aimed to predict V1 spiking activity (MUAe) from EEG recordings in a behaving macaque. We implemented two major methodological advancements: 1) using flickering stimuli to elicit high-SNR SSVEP responses in EEG, and 2) developing a comprehensive model to predict V1 MUAe using spectrotemporal features of EEG signal. We found that V1 spiking activity could be predicted from EEG, particularly during trials with flickering stimuli, both at the trial-averaged and single-trial levels. Interestingly, model performance improved further when the phase of the flickering stimulus frequency was included as an additional predictor. Subsequent analysis showed that EEG-to-MUAe prediction accuracy was higher for superficial than deep cortical layers.

### Limitations

Although we recorded multiple cortical (n=32) and EEG (n=3) electrodes across 14 sessions, with an average of 308 trials per session, our study relies on data from only a single monkey, which limits the interpretation of our findings in a few ways. First, we could not empirically verify whether the results replicate in a second monkey using a similar recording setup, a standard approach in non-human primate electrophysiology research. Second, while intracranial electrode positions cover a 3D space in V1, their positions remain fixed within V1. Because neurons in V1 exhibit temporal-frequency tuning (Foster et al., 1985; Yu et al., 2010; Zheng et al., 2007), the stimulus frequency-dependent effects – such as the stronger EEG-MUAe relationship for 10 Hz compared to other stimulus conditions – may be influenced by the specific, idiosyncratic placement of electrodes and may not generalize across the entire V1.

Therefore, in the discussion, we focus primarily on findings less likely to be limited by electrode placement. Specifically, we discuss the differences between flickering and non-flickering stimuli, as well as the effects related to model parameters that are independent of stimulus frequency. In addition, because our prediction model (Eq. 1) is designed to estimate the spiking activity envelope (MUAe), which reflects the aggregated spiking activity of multiple local neurons, its performance may be suboptimal for predicting single-unit spiking activity.

### EEG and spiking activity are related in a frequency dependent manner

Studies investigating simultaneously recorded EEG and spikes in anesthetized animals date back to the 1960s. Earlier studies involving invasive recordings from the epidural surface and spiking signals from the visual cortex in anesthetized animals showed that the majority of recorded cells fire faster during positive surface potential and fire less during negative surface potentials, both under spontaneous and visual flash stimulus conditions (Fromm & Bond, 1964; Fromm & William Bond, 1967). A series of studies by Schroeder and colleagues reported a detailed investigation of visually evoked potentials recorded from the epidural surface and underlying spiking/LFP signals in anaesthetized macaques. They showed that different components of surface potentials are contributed by the spiking activity of different visual processing areas. For example, early components (<80 ms) of evoked potentials on the cortical surface indicate V1 processing and late components (100-200 ms) indicate visual processing from extrastriate areas, such as V4 (Givre et al., 1994; Schroeder et al., 1991).

A few studies in the last two decades investigated a more detailed EEG-spike relationship using less- or non-invasive methods in behaving animals. The direct correlation between non-invasive scalp EEG and spiking activity in V4 was observed to be very weak in macaque monkeys (Snyder & Smith, 2015). However, EEG measured on the bone surface showed a clearer relationship with V1 spiking activity (Whittingstall & Logothetis, 2009). Particularly, the EEG-spike relationship was observed to be frequency dependent during naturalistic visual stimulus, where the phase of low-frequency (delta/theta) and amplitude of high-frequency (gamma) bands are better related to V1 spikes than mid-frequency bands (alpha/beta). This frequency-specific EEG to V1 spike relationship also extends to LFPs (Rasch et al., 2008).

We extend these earlier findings to non-invasive EEG, particularly by evaluating the relationship between V1 spiking signal and non-invasive occipital EEG in behaving macaques. Similar to previous LFP and invasive EEG findings, we found that EEG frequency bands are differentially related to V1 spiking activity. For the spontaneous condition (0 Hz stimulus) at a single-trial level, which can directly be compared with previous studies (Rasch et al., 2008; Whittingstall & Logothetis, 2009), we found that delta and gamma bands of EEG were correlated with MUAe (Supp. Fig. 1). However, we also observed that the alpha band shows a negative correlation, which was not observed in previous studies. We attribute this difference to the difference in the animal state and visual stimuli. A simple fixation task with no varying stimulus can be less engaging compared to the naturalistic movie-viewing paradigm used in the previous studies (Rasch et al., 2008; Whittingstall & Logothetis, 2009), hence impacting alpha amplitude and its correlation with spikes (Klimesch, 1999).

### EEG-spike relationship improves for visual stimulus eliciting steady-state responses

Repetitive visual stimuli can elicit steady-state responses in the visual cortex, characterized by high SNR (Norcia et al., 2015). Due to high SNR, these responses can easily be observed in non-invasive EEG, making them a popular choice in human neuroscience. We leveraged high-SNR responses of repetitive visual stimulus to relate spiking activity in V1 with non-invasive EEG. As expected, SSVEP responses were easily detectable in monkey EEG and reflected better EEG-spike correlation for all flickering stimulus conditions relative to spontaneous condition (Fig. 3, Fig. 4a). Furthermore, when comparing among flickering conditions, we observed that the EEG-spike relationship further depended on the stimulus frequency (Fig. 3), where relationship of spikes with each EEG band varies depending on the stimulus frequency used (Fig. 4a).

These results conform to the broader idea that visual stimuli influence the EEG-spike relationship (Snyder & Smith, 2015), and help extend and interpret the previous observations made by Whittingstall & Logothetis, (2009). They showed that delta and gamma frequency bands are better correlated with spiking activity in monkeys engaged in a naturalistic movie-viewing task, similar to our findings for single-trial spontaneous conditions (Supp. Fig. 1).

However, their observed delta-gamma relationship with spikes may originate from frequency components specific to the naturalistic movie paradigm, which elicit visual responses in the delta frequency range (3-4 Hz), as discussed in detail by Mazzoni et al., 2010. In contrast, our paradigm controlled stimulus frequencies experimentally, affirming that the EEG-spike relationship depends specifically on the stimulus frequency. Thus, we extend prior observations, suggesting that the EEG-spike relationship is contingent upon the EEG response frequencies, which can be defined by the stimulus frequencies.

### SSVEP stimuli improve EEG to spike predictions

Previous studies have shown that, during spontaneous or naturalistic stimulus conditions, delta and gamma bands in EEG can predict V1 spiking activity from both bone-surface EEG and LFP (Rasch et al., 2008; Whittingstall & Logothetis, 2009). Our findings indicate that model performance in a 0 Hz spontaneous condition was poor, likely due to the low SNR of non-invasive EEG. However, the model performed well for flickering conditions, particularly with trial-averaged signals. Although model performance with single-trial signals was generally weak, it was better for flickering stimuli compared to spontaneous conditions (Fig. 6b). This suggests that sinusoidal visual stimuli, which periodically drive the visual cortex and generate high SNR SSVEP responses (Norcia et al., 2015), enhance EEG-MUAe prediction accuracy. We speculate that visual stimuli entraining cortical neurons to an external rhythm enhance the synchronous firing of these neurons. This synchronized activity likely contributes to the EEG signal (Musall et al., 2014; Snyder et al., 2015), rendering it more predictable than the non-synchronized activity observed during spontaneous conditions.

Supporting the notion that periodic external stimuli entrain cortical neurons, Mazzoni et al. (2010) conducted a biophysically-realistic modeling study suggesting that periodic stimulation in the lateral geniculate nucleus (LGN) tunes V1 neuronal activity to the same rhythm, thereby influencing the EEG-spike relationship (Mazzoni et al., 2010). They hypothesized that low-frequency fluctuations (delta/theta) reflect shifts in cortical excitability, defining the activity of local excitatory-inhibitory (E-I) loops, which results in high-frequency activity (gamma). This combination of distinct information in low- and high-frequency bands can be used to estimate local spiking activity. Notably, Mazzoni et al. (2010) proposed that if slow fluctuations in V1 are induced externally (e.g., through LGN stimulation), this induced rhythm could enhance EEG-to-spike estimations.

To test this hypothesis, we induced rhythmic activity in V1 using visual flicker at a specific frequency rather than directly stimulating the LGN through electrical or optical methods. We based our approach on the established fact that visual information from the retina is primarily relayed to the LGN and subsequently to V1; LFP spectra in our data also show that V1 responses followed the visual flickering rhythm (not shown). Consistent with Mazzoni et al.’s hypothesis, we found that our model incorporating the stimulus phase outperformed the model without it, particularly well for single-trial estimations.

### EEG is better related to spikes at the superficial than deep cortical layers

Neocortical regions are organized in a laminar fashion, reflecting different cytoarchitectures across layers. We investigated if EEG-to-spike predictions reflect any difference across cortical layers and found that MUAe at electrode sites in shallow layers are better predicted than in deep layers (Fig. 8). While one might assume that EEG primarily reflects signals from superficial cortical layers due to volume conduction, the relatively small distance (∼500 µm) between shallow and deep layers within a cortical column suggests that volume conduction alone does not sufficiently explain why MUAe in superficial layers are better predicted with EEG signals.

Another potential explanation is that different oscillatory bands reflect distinct patterns of activity across cortical layers. The local counterpart of EEG, the local field potential (LFP), has been shown to vary along cortical layers in different regions. Notably, gamma activity displays higher relative power in superficial layers, while alpha and beta bands exhibit greater relative power in deeper layers (Mendoza-Halliday et al., 2024). Since gamma activity is closely associated with spiking activity in the visual cortex (Ray et al., 2008; Ray & Maunsell, 2011), it is reasonable to hypothesize that spikes in superficial layers may be more accurately predicted by EEG gamma activity than those in deeper layers. Our findings support this hypothesis, as we observed that spikes were better predicted in superficial layers than deep ones. To our knowledge, this study is the first to provide empirical evidence that EEG correlates more strongly with activity in superficial layers than in deep layers.

### Implications for human EEG and non-invasive brain-machine interfaces (BMI)

In this non-human primate study, we employed non-invasive EEG, the standard method for recording EEG in humans. This approach facilitates translation of our findings in human EEG and is especially valuable for advancing EEG-based non-invasive brain-machine interfaces (BMI). Under spontaneous conditions, the correlation between EEG and spiking activity is weak (Fig. 2, 3, & 4a; Snyder & Smith, 2014), making spike predictions from EEG challenging.

However, as discussed above, incorporating a broader array of EEG spectrotemporal features and SSVEP stimuli can improve this accuracy. Based on these improvements, our study offers three significant insights that are directly applicable to human EEG, paving the way for more effective non-invasive decoding of spike dynamics and associated behaviors.

First, given that EEG frequency bands are differentially related to spike dynamics (Fig. 4; Whittingstall & Logothetis, 2009), we recommend utilizing band-limited EEG rather than full-band EEG for spiking and behavioral decoding. Second, the model shows that amplitude, phase, and phase-amplitude coupling variables of EEG bands all are informative in predicting spiking signals (Supp. Fig. 2). Interestingly, phase and coupling variables were more informative than amplitude variables, particularly for single-trial estimations, indicating the potential use of phase and coupling variables in real-time BMI applications. Third, our observations that visual flicker eliciting SSVEP responses improves EEG-to-spike predictions suggests that using stimuli with periodic fluctuations may be a useful approach to improve prediction accuracy. This strategy could be leveraged in other sensory modalities and highlights the potential of stimulus-driven rhythmic stimulation to optimize neural decoding.

The improvement in EEG-to-spike predictions achieved through periodic flickering stimuli also suggests the potential for using frequency-tagging methods to investigate various cognitive functions non-invasively. Frequency tagging involves associating task-relevant cues with non-task-relevant flickering stimuli to elicit high SNR SSVEP responses in EEG (Norcia et al., 2015; Vialatte et al., 2010; Zhu et al., 2010). Our findings suggest that analyzing the spectrotemporal features of EEG responses to frequency-tagged stimuli, could result in a deeper understanding of the neural basis of complex cognitive functions in a non-invasive manner (Arora et al., 2024; Ladouce & Dehais, 2024; Peterson et al., 2014; Zhigalov et al., 2019). Future studies are needed to test this hypothesis.

---

**Supp. Fig. 1.**
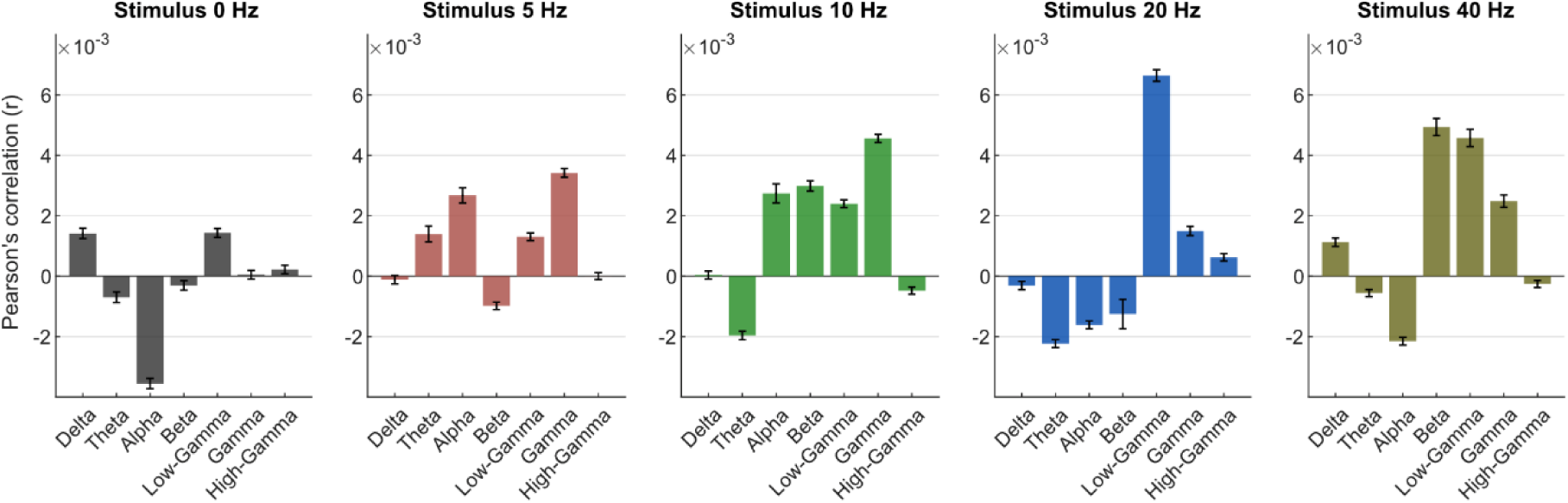
Correlation between MUAe and EEG-bands calculated at a single-trial level. All the conventions are same as in Fig. 4

**Supp. Fig. 2.**
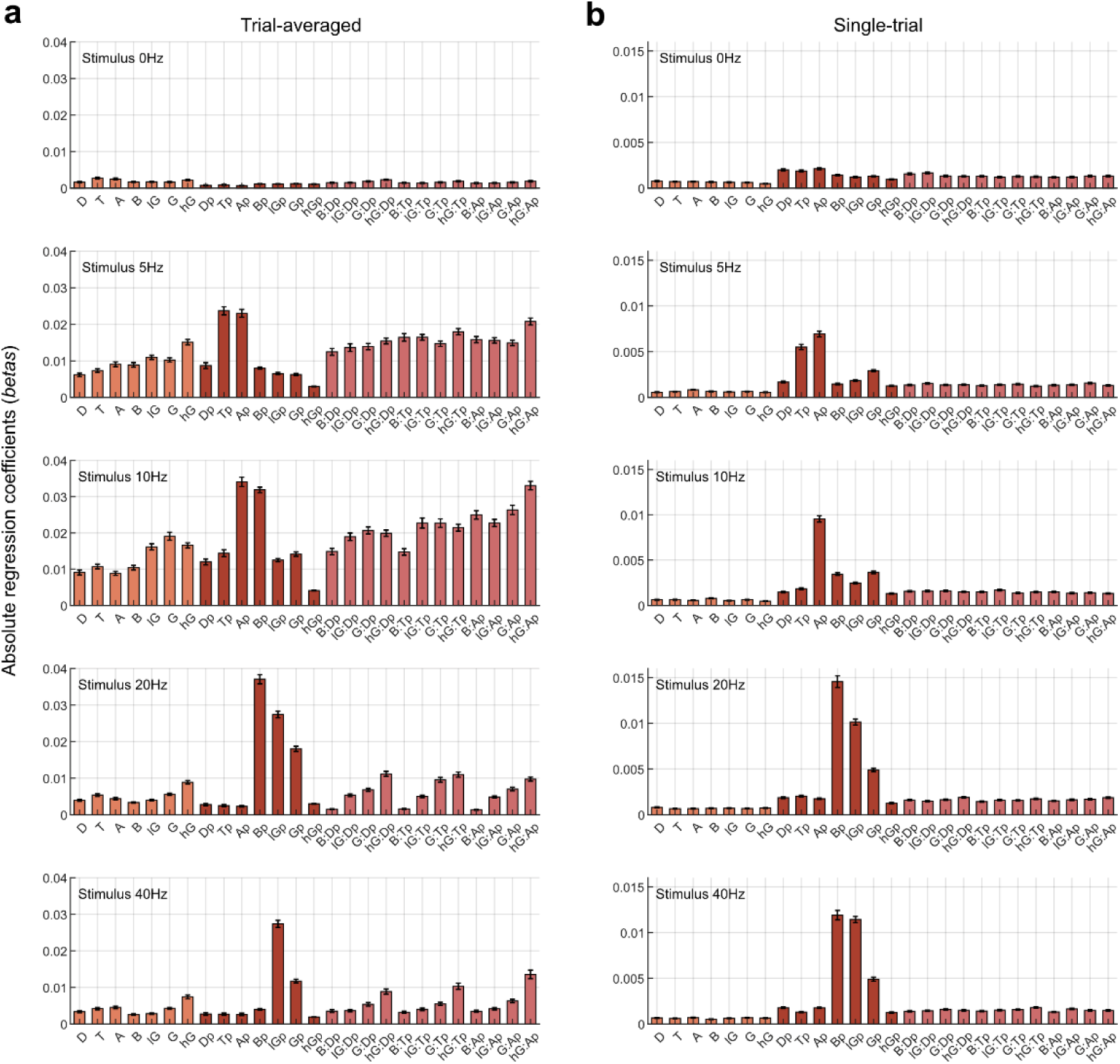
Regression coefficients of all variables of the prediction model (Eq. 1) calculated using (a) trial-averaged and (b) single-trial signals, shown separately for each stimulus condition. Each bar indicates the mean regression coefficient across all EEG-MUAe electrode pairs recorded over sessions; error bars indicate SEM. Amplitude variables are shown by single capital letters (e.g., D indicates the amplitude of EEG delta band); D: delta, T: theta, A: alpha, B: beta, lG: low-gamma, G: gamma, hG: high-gamma. Phase variables are indicated by a capital letter with ‘p’ as a suffix (e.g., Dp indicates the phase of EEG delta band). Interaction terms are indicated by a colon (:) between amplitude and phase variables (e.g., B:Dp indicates interaction between beta amplitude and delta phase variables).

## Notes

### Competing Interest Statement

The authors have declared no competing interest.

### Funding

- National Eye Institute

## REFERENCES

Alonso-Prieto, E., Belle, G. V., Liu-Shuang, J., Norcia, A. M., & Rossion, B. (2013). The 6 Hz fundamental stimulation frequency rate for individual face discrimination in the right occipito-temporal cortex. Neuropsychologia, 51(13), 2863–2875. 10.1016/j.neuropsychologia.2013.08.018

Arora, K., Gayet, S., Kenemans, J. L., Stigchel, S. V. der, & Chota, S. (2024). Rapid Invisible Frequency Tagging (RIFT) in a novel setup with EEG (p. 2024.02.01.578462). bioRxiv. 10.1101/2024.02.01.578462

Bastos, A. M., Loonis, R., Kornblith, S., Lundqvist, M., & Miller, E. K. (2018). Laminar recordings in frontal cortex suggest distinct layers for maintenance and control of working memory. Proceedings of the National Academy of Sciences, 115(5), 1117–1122. 10.1073/pnas.1710323115

Biasiucci, A., Franceschiello, B., & Murray, M. M. (2019). Electroencephalography. Current Biology, 29(3), R80–R85. 10.1016/j.cub.2018.11.052

Birbaumer, N., Murguialday, A. R., & Cohen, L. (2008). Brain–computer interface in paralysis. Current Opinion in Neurology, 21(6), 634. 10.1097/WCO.0b013e328315ee2d

Brainard, D. H. (1997). The Psychophysics Toolbox. Spatial Vision, 10(4), 433–436.

Canolty, R. T., & Knight, R. T. (2010). The functional role of cross-frequency coupling. Trends in Cognitive Sciences, 14(11), 506–515. 10.1016/j.tics.2010.09.001

Fell, J., & Axmacher, N. (2011). The role of phase synchronization in memory processes. Nature Reviews Neuroscience, 12(2), 105–118. 10.1038/nrn2979

Fisher, R. S., Harding, G., Erba, G., Barkley, G. L., & Wilkins, A. (2005). Photic- and Pattern-induced Seizures: A Review for the Epilepsy Foundation of America Working Group. Epilepsia, 46(9), 1426–1441. 10.1111/j.1528-1167.2005.31405.x

Foster, K. H., Gaska, J. P., Nagler, M., & Pollen, D. A. (1985). Spatial and temporal frequency selectivity of neurones in visual cortical areas V1 and V2 of the macaque monkey. The Journal of Physiology, 365(1), 331–363. 10.1113/jphysiol.1985.sp015776

Friston, K. J. (2009). Modalities, Modes, and Models in Functional Neuroimaging. Science, 326(5951), 399–403. 10.1126/science.1174521

Fromm, G. H., & Bond, H. W. (1964). Slow changes in the electrocorticogram and the activity of cortical neurons. Electroencephalography and Clinical Neurophysiology, 17(5), 520–523. 10.1016/0013-4694(64)90182-8

Fromm, G. H., & William Bond, H. (1967). The relationship between neuron activity and cortical steady potentials. Electroencephalography and Clinical Neurophysiology, 22(2), 159–166. 10.1016/0013-4694(67)90156-3

Givre, S. J., Schroeder, C. E., & Arezzo, J. C. (1994). Contribution of extrastriate area V4 to the surface-recorded flash VEP in the awake macaque. Vision Research, 34(4), 415–428. 10.1016/0042-6989(94)90156-2

Hastie, T., Tibshirani, R., & Wainwright, M. (2015). Statistical Learning with Sparsity: The Lasso and Generalizations. Chapman and Hall/CRC. 10.1201/b18401

Hülsemann, M. J., Naumann, E., & Rasch, B. (2019). Quantification of Phase-Amplitude Coupling in Neuronal Oscillations: Comparison of Phase-Locking Value, Mean Vector Length, Modulation Index, and Generalized-Linear-Modeling-Cross-Frequency-Coupling. Frontiers in Neuroscience, 13. 10.3389/fnins.2019.00573

Jiao, B., Li, R., Zhou, H., Qing, K., Liu, H., Pan, H., Lei, Y., Fu, W., Wang, X., Xiao, X., Liu, X., Yang, Q., Liao, X., Zhou, Y., Fang, L., Dong, Y., Yang, Y., Jiang, H., Huang, S., & Shen, L. (2023). Neural biomarker diagnosis and prediction to mild cognitive impairment and Alzheimer’s disease using EEG technology. Alzheimer’s Research & Therapy, 15(1), 32. 10.1186/s13195-023-01181-1

Klimesch, W. (1999). EEG alpha and theta oscillations reflect cognitive and memory performance: A review and analysis. Brain Research Reviews, 29(2), 169–195. 10.1016/S0165-0173(98)00056-3

Krekelberg, B., Morris, A., Cloherty, S., Koster, T., Duijnhouwer, J., Shimaoka, D., Ly, A., Hagan, M., nicholasprice, Nicroburst, Zavitz, E., & jamesmcfadyen. (2022). Neurostim: A MATLAB toolbox to run neuroscience experiments. [Computer software]. Zenodo. 10.5281/zenodo.7006340

Ladouce, S., Darmet, L., Torre Tresols, J. J., Velut, S., Ferraro, G., & Dehais, F. (2022). Improving user experience of SSVEP BCI through low amplitude depth and high frequency stimuli design. Scientific Reports, 12(1), Article 1. 10.1038/s41598-022-12733-0

Ladouce, S., & Dehais, F. (2024). Frequency tagging of spatial attention using periliminal flickers. Imaging Neuroscience, 2, 1–17. 10.1162/imag_a_00223

Lu, H.-Y., Lorenc, E. S., Zhu, H., Kilmarx, J., Sulzer, J., Xie, C., Tobler, P. N., Watrous, A. J., Orsborn, A. L., Lewis-Peacock, J., & Santacruz, S. R. (2021). Multi-scale neural decoding and analysis. Journal of Neural Engineering, 18(4), 045013. 10.1088/1741-2552/ac160f

Mazzoni, A., Whittingstall, K., Brunel, N., Logothetis, N. K., & Panzeri, S. (2010). Understanding the relationships between spike rate and delta/gamma frequency bands of LFPs and EEGs using a local cortical network model. NeuroImage, 52(3), Article 3. 10.1016/j.neuroimage.2009.12.040

Mendoza-Halliday, D., Major, A. J., Lee, N., Lichtenfeld, M. J., Carlson, B., Mitchell, B., Meng, P. D., Xiong, Y. (Sophy), Westerberg, J. A., Jia, X., Johnston, K. D., Selvanayagam, J., Everling, S., Maier, A., Desimone, R., Miller, E. K., & Bastos, A. M. (2024). A ubiquitous spectrolaminar motif of local field potential power across the primate cortex. Nature Neuroscience, 1–14. 10.1038/s41593-023-01554-7

Mitzdorf, U. (1985). Current source-density method and application in cat cerebral cortex: Investigation of evoked potentials and EEG phenomena. Physiological Reviews, 65(1), 37–100. 10.1152/physrev.1985.65.1.37

Musall, S., von Pföstl, V., Rauch, A., Logothetis, N. K., & Whittingstall, K. (2014). Effects of Neural Synchrony on Surface EEG. Cerebral Cortex, 24(4), Article 4. 10.1093/cercor/bhs389

Norcia, A. M., Appelbaum, L. G., Ales, J. M., Cottereau, B. R., & Rossion, B. (2015). The steady-state visual evoked potential in vision research: A review. Journal of Vision, 15(6), 4. 10.1167/15.6.4

Nunez, P. L., & Srinivasan, R. (2006). Electric Fields of the Brain: The Neurophysics of EEG. Oxford University Press.

Pelli, D. G. (1997). The VideoToolbox software for visual psychophysics: Transforming numbers into movies. Spatial Vision, 10(4), 437–442.

Peterson, D. J., Gurariy, G., Dimotsantos, G. G., Arciniega, H., Berryhill, M. E., & Caplovitz, G. P. (2014). The steady-state visual evoked potential reveals neural correlates of the items encoded into visual working memory. Neuropsychologia, 63, 145–153. 10.1016/j.neuropsychologia.2014.08.020

Rasch, M. J., Gretton, A., Murayama, Y., Maass, W., & Logothetis, N. K. (2008). Inferring Spike Trains From Local Field Potentials. Journal of Neurophysiology, 99(3), 1461–1476. 10.1152/jn.00919.2007

Ray, S., Crone, N. E., Niebur, E., Franaszczuk, P. J., & Hsiao, S. S. (2008). Neural Correlates of High-Gamma Oscillations (60–200 Hz) in Macaque Local Field Potentials and Their Potential Implications in Electrocorticography. Journal of Neuroscience, 28(45), Article 45. 10.1523/JNEUROSCI.2848-08.2008

Ray, S., & Maunsell, J. H. R. (2011). Different Origins of Gamma Rhythm and High-Gamma Activity in Macaque Visual Cortex. PLOS Biology, 9(4), Article 4. 10.1371/journal.pbio.1000610

Schroeder, C. E., Tenke, C. E., Givre, S. J., Arezzo, J. C., & Vaughan, H. G. (1991). Striate cortical contribution to the surface-recorded pattern-reversal vep in the alert monkey. Vision Research, 31(7), 1143–1157. 10.1016/0042-6989(91)90040-C

Snyder, A. C., Morais, M. J., Willis, C. M., & Smith, M. A. (2015). Global network influences on local functional connectivity. Nature Neuroscience, 18(5), 736–743. 10.1038/nn.3979

Snyder, A. C., & Smith, M. A. (2015). Stimulus-dependent spiking relationships with the EEG. Journal of Neurophysiology, 114(3), 1468–1482. 10.1152/jn.00427.2015

Supèr, H., & Roelfsema, P. R. (2005). Chronic multiunit recordings in behaving animals: Advantages and limitations. In Progress in Brain Research (Vol. 147, pp. 263–282). Elsevier. 10.1016/S0079-6123(04)47020-4

Tenke, C. E., Schroeder, C. E., Arezzo, J. C., & Vaughan, H. G. (1993). Interpretation of high-resolution current source density profiles: A simulation of sublaminar contributions to the visual evoked potential. Experimental Brain Research, 94(2), 183–192. 10.1007/BF00230286

Tort, A. B. L., Kramer, M. A., Thorn, C., Gibson, D. J., Kubota, Y., Graybiel, A. M., & Kopell, N. J. (2008). Dynamic cross-frequency couplings of local field potential oscillations in rat striatum and hippocampus during performance of a T-maze task. Proceedings of the National Academy of Sciences, 105(51), 20517–20522. 10.1073/pnas.0810524105

Vialatte, F.-B., Maurice, M., Dauwels, J., & Cichocki, A. (2010). Steady-state visually evoked potentials: Focus on essential paradigms and future perspectives. Progress in Neurobiology, 90(4), 418–438. 10.1016/j.pneurobio.2009.11.005

Whittingstall, K., & Logothetis, N. K. (2009). Frequency-Band Coupling in Surface EEG Reflects Spiking Activity in Monkey Visual Cortex. Neuron, 64(2), 281–289. 10.1016/j.neuron.2009.08.016

Xing, D., Yeh, C.-I., Burns, S., & Shapley, R. M. (2012). Laminar analysis of visually evoked activity in the primary visual cortex. Proceedings of the National Academy of Sciences, 109(34), 13871–13876. 10.1073/pnas.1201478109

Yu, H.-H., Verma, R., Yang, Y., Tibballs, H. A., Lui, L. L., Reser, D. H., & Rosa, M. G. P. (2010). Spatial and temporal frequency tuning in striate cortex: Functional uniformity and specializations related to receptive field eccentricity. European Journal of Neuroscience, 31(6), 1043–1062. 10.1111/j.1460-9568.2010.07118.x

Zheng, J., Zhang, B., Bi, H., Maruko, I., Watanabe, I., Nakatsuka, C., Smith, E. L., & Chino, Y. M. (2007). Development of Temporal Response Properties and Contrast Sensitivity of V1 and V2 Neurons in Macaque Monkeys. Journal of Neurophysiology, 97(6), 3905–3916. 10.1152/jn.01320.2006

Zhigalov, A., Herring, J. D., Herpers, J., Bergmann, T. O., & Jensen, O. (2019). Probing cortical excitability using rapid frequency tagging. NeuroImage, 195, 59–66. 10.1016/j.neuroimage.2019.03.056

Zhu, D., Bieger, J., Garcia Molina, G., & Aarts, R. M. (2010). A Survey of Stimulation Methods Used in SSVEP-Based BCIs. Computational Intelligence and Neuroscience, 2010(1), 702357. 10.1155/2010/702357

